# Development of ferret immune repertoire reference resources and single-cell-based high-throughput profiling assays

**DOI:** 10.1101/2025.02.05.636682

**Authors:** Evan S. Walsh, Kui Yang, Tammy S. Tollison, Sujatha Seenu, Nicole Adams, Guilhem Zeitoun, Ifigeneia Sideri, Geraldine Folch, Hayden N. Brochu, Hsuan Chou, Sofia Kossida, Ian A. York, Xinxia Peng

**Author notes:** **Contact info**, Correspondence. These authors contributed equally (Evan S. Walsh, Kui Yang). 1300 Rancho Conejo Blvd, Thousand Oaks, CA 91320.

## Abstract

Domestic ferrets (*Mustela putorius furo*) are important for modeling human respiratory diseases. However, ferret B and T cell receptors have not been completely identified or annotated, limiting immune repertoire studies. Here we performed long read transcriptome sequencing of ferret splenocyte and lymph node samples to obtain over 120,000 high-quality full-length immunoglobin (Ig) and T cell receptor (TCR) transcripts. We constructed a complete reference set of the constant regions of ferret Ig and TCR isotypes and chain types. We also systematically annotated germline Ig and TCR variable (V), diversity (D), joining (J), and constant (C) genes on a recent ferret reference genome assembly. We designed new ferret-specific immune repertoire profiling assays by targeting positions in constant regions without allelic diversity across 11 ferret genome assemblies, and experimentally validated them using a commercially compatible single-cell-based platform. These improved resources and assays will enable future studies to fully capture ferret immune repertoire diversity.

## INTRODUCTION

Domestic ferrets (*Mustela putorius furo*) are an increasingly common model organism to study human respiratory diseases. For example, ferrets are routinely used in studies involving influenza viruses due to their ability to be infected with human strains ^1, 2^, allowing vaccine ^3^ and therapeutic ^4, 5^ efficacy studies. Ferrets are also useful models for cystic fibrosis ^6^, and more recently ferrets have been used as a model organism of SARS-CoV-2 infection ^7–9^. Despite these important roles in translational medicine, ferret infectious disease research is limited due to a lack of resources, including incomplete and missing immunogenetic resources. Developing complete and accurate genomic resources, especially for the immune system, is imperative for effective translational interpretation.

B and T cell receptor molecules serve the important role of recognizing and binding to foreign antigens (Ag) from infectious agents. The ability of these receptors to recognize foreign antigens is in part due to their generation and structure. Immunoglobulins (Igs) and T cell receptors (TCRs) have two domains: a constant (C) region and a variable (V) region. The V region is comprised of a variable (V), joining (J), and in some cases, a diversity (D) gene. These genes are duplicated in several large loci in the genome and are somatically rearranged before transcription to establish a diverse repertoire of both receptor molecules and antigen binding sites. Following rearrangement, humans may have up to 10^13^ and 10^18^ unique Ig and TCR V-region domains within an individual, respectively ^10^. The genetic diversity of Ig and TCR molecules makes their correct assembly, annotation, and validation a major technical challenge for standard high-throughput short-read based approaches, as illustrated by earlier artificial chromosome cloning based studies ^11, 12^ and more recent ones based on long-read sequencing ^13, 14^.

Immune repertoire studies are rapidly becoming a mainstream tool for understanding immunity to infectious and autoimmune diseases and cancers. In these studies, the Ig and TCR repertoires of B and T lymphocytes are profiled, usually via high-throughput sequencing methods (Rep-seq). Currently only humans, mice, and some non-human primates (rhesus monkey ^15^, lowland gorilla ^16^) have relatively complete germline Ig and TCR annotations and established assays needed to perform and analyze high-throughput immune-profiling analyses. While progresses such as ^17–19^ have been made to perform and analyze high-throughput immune-profiling analyses in other species, ferrets remain lacking in this area of study, making the analysis and interpretation of immune repertoire studies challenging.

Some efforts have been made to close the gap in ferret immunogenetics and annotate these complex Ig and TCR genes. Recently, Wong et al. ^20^ predicted V-, D-, J-, and C-region genes from Ig loci and subsequently designed a single-cell multiplex PCR (MPCR) experiment for sequence validation. Additionally, Gerritsen et al. ^21^ used 5’ rapid amplification of cDNA ends (RACE) to annotate the ferret T cell receptor B (TRB) locus. These were the only ferret sequences available in the international ImMunoGeneTics information system [IMGT] database^22^. The most recent annotation of the domestic ferret genome was the NCBI RefSeq annotation release 102 on the NCBI RefSeq genome assembly GCF_011764305 ^23^ (the corresponding Submitted GenBank assembly was GCA_011764305.2 which was generated using both Illumina short read and PacBio long read technologies ^24^). This RefSeq annotation reportedly contains 142 Ig and TCR genes ^25^, which are computational predictions based on orthologous genes in similar species and have yet to be validated. Despite these recent advances, the design of ferret-specific assays for profiling Ig and TCR repertoires still rely on very limited resources.

In this study, we aimed to systematically identify and annotate ferret Ig and TCR sequences, including the C, V, D, and J genes of each, by extending previously established strategies ^18, 26^ and combining multiple approaches. This allowed us to construct the first complete reference set of ferret Ig and TCR C-region genes, covering all ferret Ig/TCR isotypes and chain types, using long read transcriptome sequencing analysis (Iso-Seq). We then utilized the Iso-Seq data, C-region reference, and recombination signal sequence (RSS) analysis to additionally identify hundreds of Ig and TCR V, D, and J loci on a recent ferret reference genome assembly. We also used this C-region reference to design new ferret-specific assays that allow for a full Ig/TCR repertoire analysis at the single cell level. We validated these new assays with ferret peripheral blood mononuclear cell (PBMC) and splenocyte samples in conjunction with the 10x Genomics Chromium system, a common commercial single-cell sequencing platform. Taken together, this study provides comprehensive reference resources and assays that are essential for ferret immunogenetics and immune repertoire profiling analysis.

## RESULTS

### Overall strategy

We started by performing Iso-Seq analysis of ferret lymph node and spleen samples, similarly as^26^ to circumvent the challenging task of correctly assembling Ig/TCR transcripts from short sequencing reads. The complete reference set of ferret Ig and TCR constant region (C-region) sequences was constructed from the obtained high-quality full-length Ig and TCR transcript sequences and the corresponding ferret C-region genes were annotated on a recent ferret reference genome assembly (NCBI accession no: GCA_011764305.2)

Guided by these ferret Ig/TCR C-region references, we selected full-length Ig and TCR transcript sequences containing C-regions from the ferret Iso-Seq data and analyzed them for V, D, and J genes. In parallel, we performed semi-automated analysis of ferret genome assemblies, using conserved features such as RSS sequences to identify and annotate putative germline genes. Germline V, D, J genes were manually reviewed, curated, and classified as functional, ORF, or pseudogenes based on the presence of conserved features, including the RSS and full-length open reading frames. Subsequently, expression of V and J genes in transcriptomes from naïve ferrets was used to further validate the classification of genes as functional; this resulted in changing the classification of several genes (mainly those with non-canonical RSS sequences; see below) to “functional”.

We then designed new ferret-specific V(D)J assays that use primers intended to target the C-regions and allow single cell level Ig/TCR repertoire analysis, similarly as in ^18, 26^. Individual ferret specific primers were experimentally validated and screened *in silico* across 11 ferret genome assemblies to account for observed variations in the ferret C-regions including allelic, isoform, and subtype. Finally, these new ferret-specific assays were evaluated by single-cell based V(D)J and transcriptome sequencing analysis of ferret PBMC and splenocyte samples on a commercial single-cell platform.

### Generation of a complete collection of reference sequences and annotations for domestic ferret Ig and TCR constant region genes

Using PacBio Iso-Seq transcriptome sequencing, we obtained over 5.9 million circular consensus sequence (CCS) reads from 11 samples sourced from either lymph nodes or splenocytes. Each CCS read represents the consensus sequence of a single transcript. About 89.75% of these CCS reads were full length (i.e., contained the 5’ cDNA primer, 3’ cDNA primer, and polyadenylation tail). Using the available IMGT annotation of the V, D, and J regions of Ig and TCR germline sequences from ferret, horse, dog, and cat as an IgBLAST database, we recovered 118,485 Ig and 4,746 TCR putative transcript sequences (2% and 0.08% of the total number of full-length CCS reads) (Table S1). The average rates of mismatch, insertion, and deletion events in the C region sequences were estimated to be 0.11%, 0.12%, 0.04% respectively, which were comparable to our previous observations ^26^.

We extracted the C-regions from these putative Ig/TCR transcript sequences and constructed C-region consensus sequences from 202 clusters of highly similar sequences. These consensus sequences represented complete cDNA sequences for the entire ferret C-regions (i.e., including the 3’ UTR). We then filtered out low quality consensus sequences and collapsed redundant consensus sequences representing the same transcript isoforms from the same genes based on their alignments to the reference ferret genome assembly sequences (GCA_011764305.2), to generate a set of non-redundant C-region consensus sequences. Further, we only included consensus sequences with complete open read frames (ORFs) and examined the corresponding splice site sequences after aligning them to the ferret reference genome assembly. These ferret C-region reference sequences and annotations were further manually curated and named based on the orthologous human C-region genes ^15, 27, 28^. Additional details are described in the Methods section.

As summarized in Table 1, Figures 1 and 2, Figure S1, and Table S2, we identified functional ferret orthologues for IGHA, IGHD, IGHE, IGHG2, IGHG4, IGHM, IGKC, IGLC, TRBC, TRDC, and TRGC. We also identified ferret orthologues for the components of the surrogate light chain, VpreB and λ5 (Table 1 and Figure 1D). We detected multiple subclasses of ferret IGHG, IGLC, TRBC, and TRGC genes, based on the alignment of consensus sequences to the reference ferret genome assembly (GCA_011764305.2). For each IGH isotype, we also identified both membrane-bound and secreted transcript splicing isoforms in the transcriptome (Table 1). We also observed apparent allelic variants (i.e. genetic variations among ferrets evidenced by the sequence differences between the transcripts we obtained in this study and the reference genome assembly) for each of the IGH isotypes data (Table 1 and Table S2 “Allelic Variants” tab). We also observed several additional alternatively spliced transcript isoforms of low frequencies in the Iso-Seq data which are not included here, as it is unclear if they were technical artifacts. Overall, ferret Ig and TCR C-region genes are present on the reference assembly in the same relative order and orientation as seen in humans (Figure 1A-C, Figure 2A-D).

**Table 1.** Summary of Ferret Ig and TCR Constant Regions. Ferret Ig and TCR constant regions and allelic variants were identified as described in the Methods. Transcriptomes comprising splenocytes from a total of 10 individual ferrets (SRA accession numbers SRR29376976, SRR29376975, SRR29376974, SRR26825672 and SRR26825671) were assessed for the number of exact matches for each sequence. See Table S2 for detailed sequence information, and Table S2 “llelic Variants” tab for sequences and counts of allelic variants. N/A: The CCS read counts of pseudogenes and surrogate light chain genes were not included.

| Subclass | Form | Number of allelic variants | CCS counts (10 ferrets) |
| --- | --- | --- | --- |
| IGHA | Secreted | 4 | 14197 |
|  | Transmembrane |  | 336 |
| IGHD | Secreted | 2 | 266 |
|  | Transmembrane |  | 466 |
| IGHE | Secreted | 3 | 5 |
|  | Transmembrane |  | 0 |
| IGHG2 | Secreted | 3 | 332500 |
|  | Transmembrane |  | 10276 |
| IGHG4 | Secreted | 2 | 28572 |
|  | Transmembrane |  | 322 |
| IGHM | Secreted | 2 | 9227 |
|  | Transmembrane |  | 7942 |
| IGKC |  | 1 | 2519 |
| IGLC1 |  | 1 | 3680 |
| IGLC2 |  | 1 | 7 |
| IGLC3 |  | 1 | 1206 |
| IGLC4 | Pseudogene | 1 | N/A |
| IGLC5 |  | 1 | 1206 |
| IGLC6 | Pseudogene | 1 | N/A |
| IGLC7 |  | 1 | 86 |
| TRAC1 |  | 1 | 33 |
| TRBC1 |  | 2 | 79 |
| TRBC2 |  | 1 | 109 |
| TRDC1 |  | 1 | 429 |
| TRGC1 | Pseudogene | 1 | N/A |
| TRGC2 | Pseudogene | 1 | N/A |
| TRGC3 |  | 1 | 43 |
| TRGC4 |  | 1 | 1 |
| TRGC5 |  | 1 | 38 |
| TRGC5A | Pseudogene | 1 | N/A |
| TRGC6 | Pseudogene | 1 | N/A |
| TRGC7 |  | 1 | 0 |
| TRGC8 |  | 1 | 0 |
| TRGC9 |  | 1 | 0 |
| VpreB |  | 1 | N/A |
| $\lambda 5$ | | 1 | N/A |

**Figure 1.**
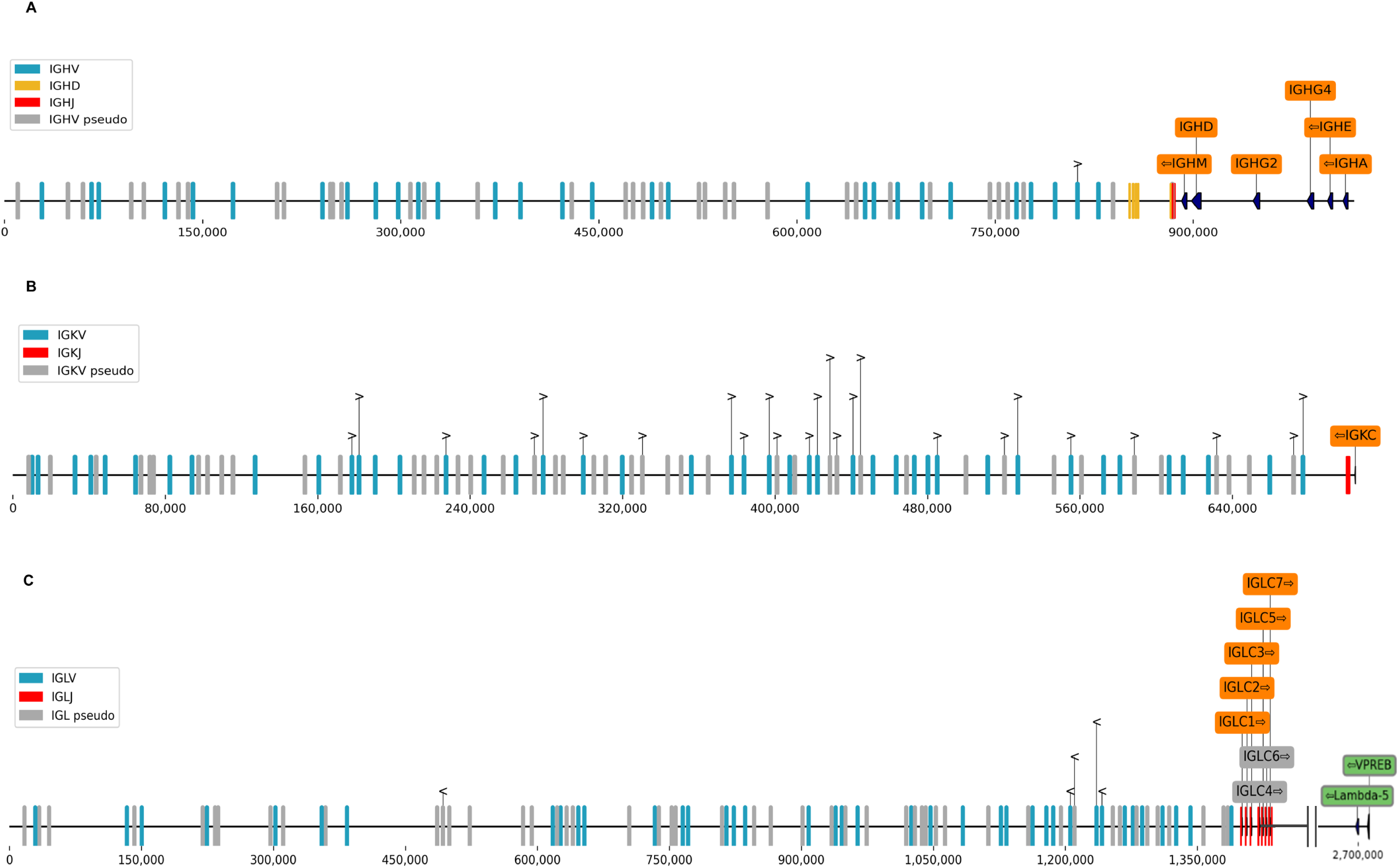
Genomic organization of ferret Ig regions. (A) Heavy chains and associated V, D, and J genes; (B) Kappa light chains and associated V and J genes; (C) Lambda light chains and associated V and J genes. Constant regions are annotated in orange, V regions are in blue, D regions are in yellow, J regions are in red, and surrogate light chain components VpreB and λ5 are shown in grey. The orientation of each gene is indicated with an arrow. Kappa annotations are on the sense strand of the reference contig but depicted on the antisense strand in (C). Genome positions relative to ferret reference genome assembly GCA_011764305.2, contigs JAADYL010000038.1 (heavy chain locus), JAADYL010000784.1 (light chain lambda locus), and JAADYL010000091.1 (light chain kappa locus) are indicated below the bar. See also Figure S1 for additional details.

**Figure 2.**
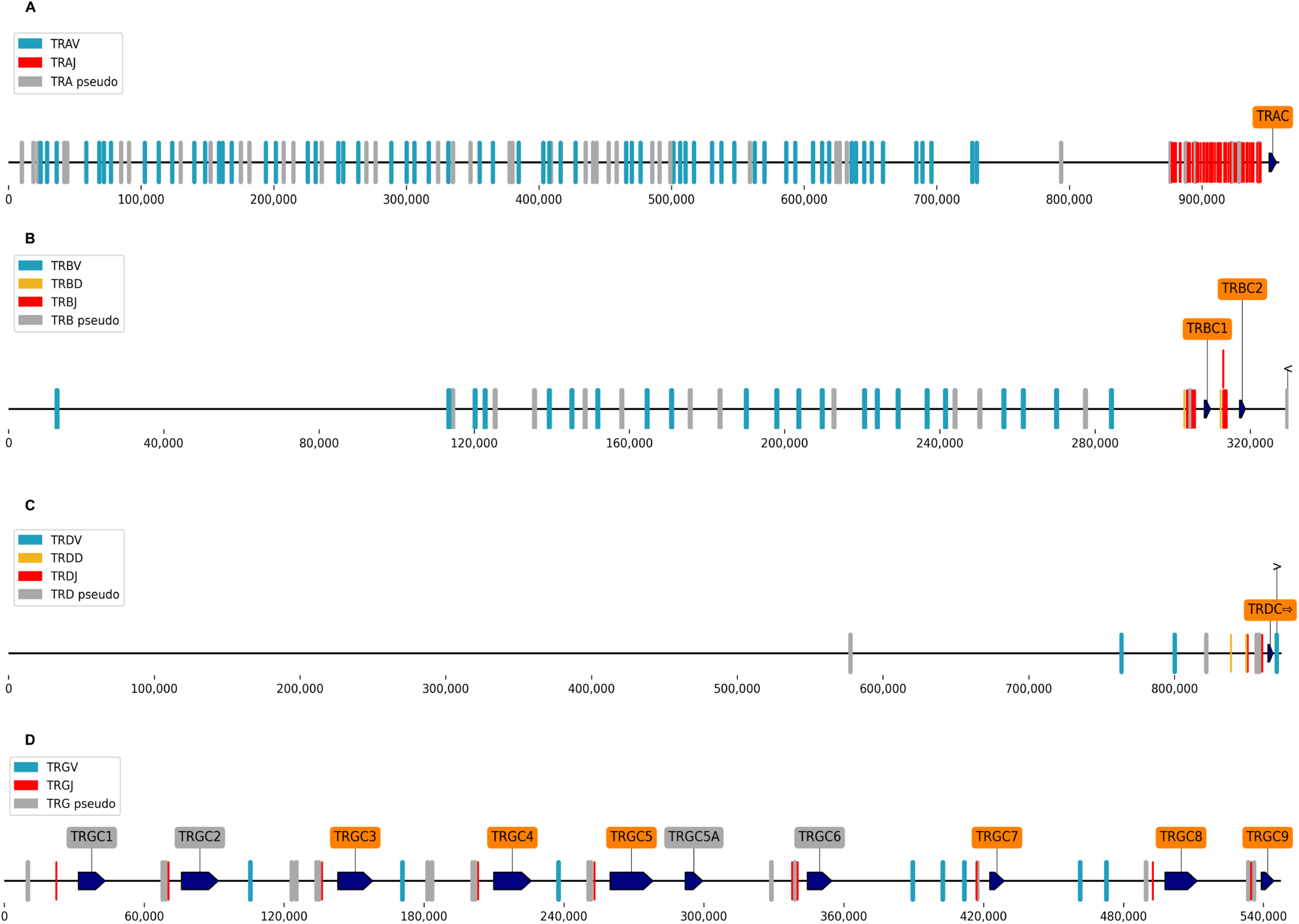
Genomic organization of ferret TCR region. (A) TRA associated V, D and J genes; (B) TRB and associated V, D, and J genes; (C) TRG and associated V and J genes; (D) TRD and associated V and J genes. Constant regions are annotated in orange, V regions are in blue, D regions are in yellow, and J regions are in red. The orientation of each gene is indicated with an arrow. Genome positions relative to ferret reference genome assembly GCA_011764305.2, contigs JAADYL010000821.1 (TCR alpha and delta chain locus), JAADYL010000772.1 (TCR beta and gamma chain locus) are indicated below the bar. See also Figure S1 for additional details.

### Generation of a reference annotation of domestic ferret Ig and TCR V, D, and J germline genes

We further identified and annotated Ig and TCR V, D, and J germline genes on the recent ferret reference genome assembly (GCA_011764305.2) using a combined analysis of the ferret Ig/TCR transcripts and genomic sequences, and manually validated and curated them based on the presence of highly conserved features such as RSS signals (see Methods and Figure S2). The number of Ig and TCR V-, D-, and J-region genes are summarized in Table 2 and their genome arrangements are shown in Figures 1 and 2 and Figure S1. Detailed annotations of Ig and TCR V, D, J genes are provided in Table S2 and Supplementary Text. A significant number of ferret Ig V genes were identified as pseudogenes or non-functional open reading frames (ORFs) (35/63 for IGH, 96/136 for IGL, and 47/87 for IGK). For TRA, TRB, TRD, and TRG, we identified 96, 34, 5, and 16 V genes (with 44, 15, 2, and 10 being non-functional), respectively (Table 2; Table S2).

**Table 2.** Summary of Ferret V, D, and J Region Counts for Ig and TCR. Ferret Ig and TCR V, D, and J regions were identified and classified as functional, pseudogenes, or ORF, as described in the Methods. In the case of IGHV, 19 provisional genes found in separate contigs (see Methods; Table S2, “IGHV Provisional” tab) are not included. See also Table S2.

| Class | Gene type | Functional | ORF | Pseudogene | Total |
| --- | --- | --- | --- | --- | --- |
| IGH | V | 28 | 1 | 34 | 63 |
|  | D | 9 | 1 | 0 | 10 |
|  | J | 6 | 1 | 1 | 8 |
| IGK | V | 40 | 3 | 44 | 87 |
|  | J | 4 | 1 | 0 | 5 |
| IGL | V | 40 | 6 | 90 | 136 |
|  | J | 7 | 1 | 0 | 8 |
| TRA | V | 52 | 9 | 35 | 96 |
|  | J | 36 | 19 | 5 | 60 |
| TRB | V | 19 | 3 | 12 | 34 |
|  | D | 2 | 0 | 0 | 2 |
|  | J | 8 | 3 | 1 | 12 |
| TRD | V | 3 | 0 | 2 | 5 |
|  | D | 2 | 0 | 0 | 2 |
|  | J | 2 | 0 | 3 | 5 |
| TRG | V | 6 | 2 | 8 | 16 |
|  | J | 4 | 6 | 11 | 21 |

Ferret germline genes were generally located to the expected contexts, based on comparisons to other species (Figures 1 and 2). For example, IGH V genes generally map upstream of the heavy chain constant regions on the same contig (Figure 1A) and IgL J genes map between each of the seven IgL gene loci (Figure 1C) ^18, 19^. Most ferret RSSs identified in this study shared the canonical heptamer (CACAGTG) and nonamer (ACAAAAACC) sequences of both mouse and human (Table S2 and Figure S3); however, abundant expression of several genes flanked by non-canonical heptamers and/or nonamers was identified (Table S2); for example, some well-expressed genes used CGATTCGGA, CCATATTGT, GTCTTTGTC, or ACTTCTTGT instead of the canonical nonamer, or CAGTGTG or CATTGTG instead of the canonical heptamer.

### Design of ferret-specific B and T cell V(D)J assays for single cell analysis

We designed ferret-specific primer sets based on the Ig and TCR constant regions identified above, using consensus sequences that match all isoforms of each receptor class. We also checked these primers against 11 ferret genome assemblies that were made public recently to reduce the potential effects of allelic variations on the amplification coverage (see Methods). Primer sets are described in Table S3.

We validated the ability of our primers to amplify each of the C-region references *in silico* (Figure S4) and their amplification specificity experimentally (Figure 3A), and compared them to reverse primers targeting C-regions of Ig and TRB genes from Wong et al. ^20^ and Gerritsen et al. ^21^, respectively (Figure S4). Across every Ig and TCR C-region gene in our reference we were able to amplify all of the corresponding C-region genes identified on the 11 ferret genome assemblies *in silico*, with the exception of the pseudogene TRGC6*01 with a truncated exon 1 sequence that does not have the inner primer binding site (Figure S4). This coverage was better than or equal to that of primers by Wong et al. ^20^ and Gerritsen et al. ^21^ across all genes (Figure S4).

**Figure 3.**
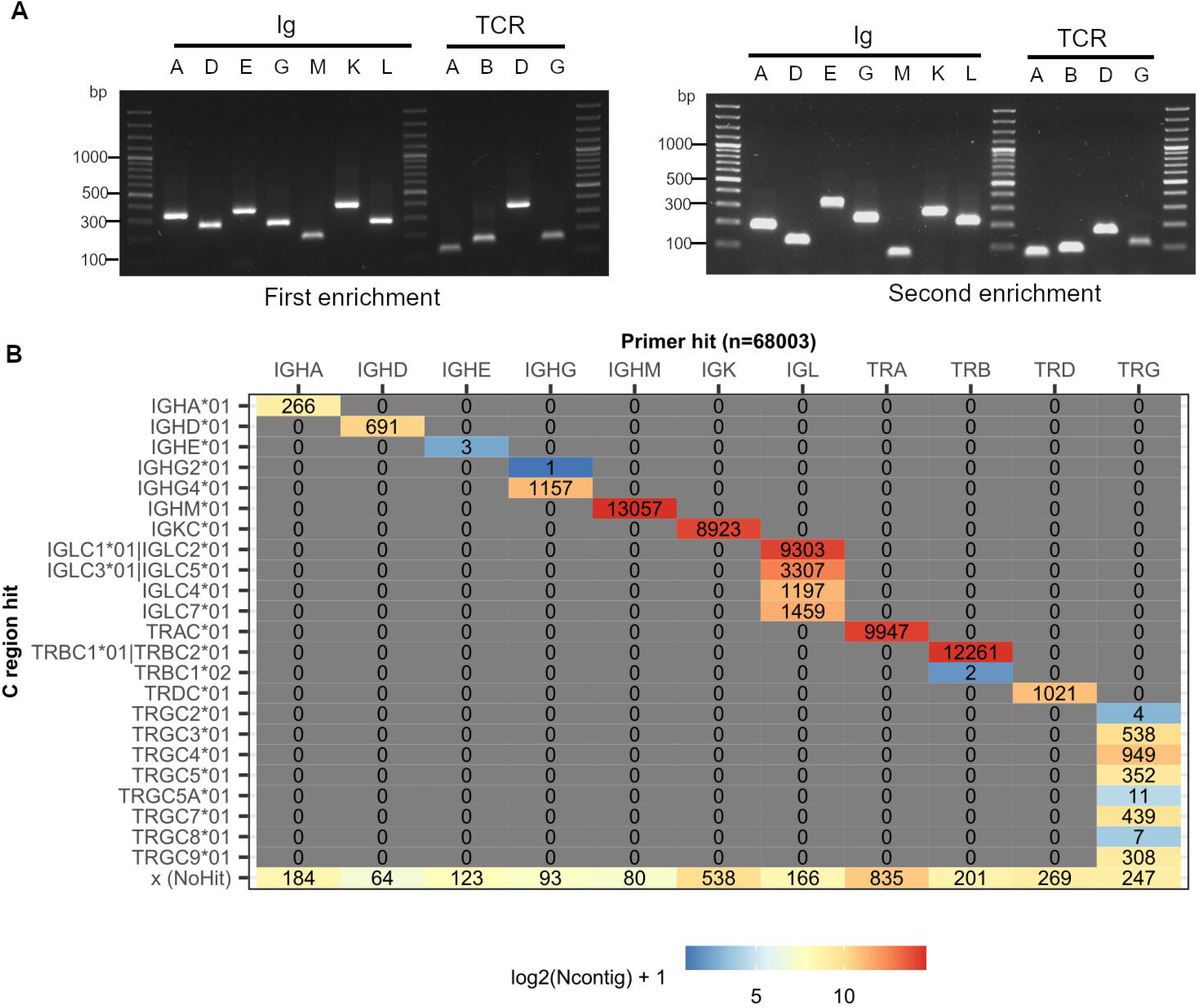
Validation of ferret-specific primers. (A) Individual ferret Ig and TCR specific primers were used to amplify the corresponding genes of the C-region genes from cDNAs isolated from a pool of three PBMC samples. Corresponding forward primers that were upstream but still within the C regions were designed solely for the purposes of this assay. PCR amplicon products of such reactions were analyzed on a 1% agarose gel to confirm the amplification specificity. Gel image on the left: primers for the first enrichment reactions. Gel image on the right: primers for the second enrichment reactions. On each gel image lanes 1, 9, and 14 are 100bp ladder. Lanes for Igs are labeled as IGHA (A), IGHD (D), IGHE (E), IGHG (G), IGHM (M), IGK (K), and IGL (L). Lanes for TRs are labeled as TRA (A), TRB (B), TRD (D), and TRG (G). (B) Recovery of ferret V(D)J transcripts using the single cell based V(D)J assays with the ferret C-region specific primers. The tiled plot indicates the total number of ferret V(D)J transcript contigs with respective ferret primer and C region matches recovered from the ferret PBMC and splenocyte samples. See also Figure S4.

### Immune repertoire sequencing of ferret PBMC and splenocyte tissue samples

We next sought to experimentally validate these ferret-specific single-cell based V(D)J assays. To this end, we applied our ferret-specific primers to ferret PBMC and splenocyte tissue samples in conjunction with the 10x Genomics based paired gene expression and immune repertoire analysis workflow, similarly as ^18^. These V(D)J assays enable the recovery of corresponding transcriptomic profiles and paired VDJ-VJ chains from single B and T cells. We sequenced over 16,000 single-cell barcodes from PBMC and splenocyte tissue samples. De novo assembly of Ig and TCR enrichment sequencing libraries resulted in 82,418 unfiltered, unannotated contigs. As shown in Figure 3B, 68,003 contigs had ferret Ig/TCR enrichment primer hits and 65,203 (95.9%) of them also had ferret Ig/TCR C-region matches which all corresponded to the primer hit, indicating our ferret Ig/TCR enrichment primers were efficient and specific. All ferret Ig/TCR isotypes and subtypes, including the weakly expressed IGHE isotype, were recovered. The small percentage of contigs with primer hits but not C-region matches tended to have lower UMI counts and were shorter (Table S3), suggesting that their corresponding transcripts might not be fully assembled *de novo* due to the low transcript abundances.

Separately we filtered single cell transcriptomic profiles for cells of healthy quality and removed cell barcodes suspected of being technical doublets (see Methods). After filtering in total we obtained 12,331 cell barcodes (8,016 for PBMC and 4,315 for splenocyte sample), 5,550 barcodes with annotated Ig VDJ and/or VJ contigs and 5,776 barcodes with TCR VDJ and/or VJ contigs. Also 810 cell barcodes had annotated TRG or TRD contigs. Interestingly, we observed a large difference between the two samples in terms of the frequencies of gamma-delta T cells, with 400 cell barcodes with TRG and/or TRD contig in the PBMC sample (8.1% of all cell barcodes with at least one TCR contig) and 410 in the splenocyte sample (49.3%).

Parallel gene expression analysis allowed us to cluster single cells and served as a ground truth to assess the pairing efficiency of our single-cell immune repertoire assay. Pairing efficiency is defined as the rate of captured B and T cells that have paired VDJ and VJ chains recovered. To integrate gene expression profiles and immune repertoires of single cells we clustered PBMC and splenocyte cells in UMAP space and examined the VDJ-VJ pairing efficiency in each cluster with at least 50 cells. Based on the detection of Ig contigs, we captured four clusters of B cells with at least 50 cells in the PBMC sample (Figure 4A & 4B, Table S4). Figure 4C shows the Ig pairing efficiencies in the ferret PBMC sample, which range from 84.8% (cluster 3) to 89.8% (cluster 5). Among three T cell clusters in the PBMC sample, the TCR pairing efficiencies were 60.7% for cluster 1 and 72.2% for cluster 2, but 41% for cluster 8 (Figure 4C). To investigate the apparently low TCR pairing efficiency for cluster 8 and to check if our Ig/TCR analysis might have missed other B and T cell clusters, we performed the cell type prediction for each cell using SingleR ^29^ (see Methods, Figure S5, and Table S5). We also examined the expression of ferret orthologs of several canonical markers for human B (CD79A, CD79B, CD19) and T (CD3D, CD3D, CD3G) cells in each of the cell clusters (Figure S6). We found that cluster 8 included two subclusters of cells, one for T and one for NK cells (Figure S5 A and B), and the TCR pairing efficiency for the T cell subcluster was 63.3% (Figure S5 C). The same cell type prediction analysis also indicated that cluster 6 was a mixture of B and T cells (Figure S5 B), but the observed Ig or TCR efficiency was extremely low due to the lack of detection of Ig/TCR contigs (Figure 4A). We compared the cell gene expression profiles of cluster 6 to those of clusters 2 (closest T cell cluster in Figure 4A) and 7 (closest B cell cluster). Functional analysis of both lists of the differentially expressed genes showed that cluster 6 was highly enriched with cell death related biological functions (Table S6), suggesting cells in cluster 6 might have failed to fully rearrange their VDJ/VJ regions, leading to programmed cell death.

**Figure 4.**
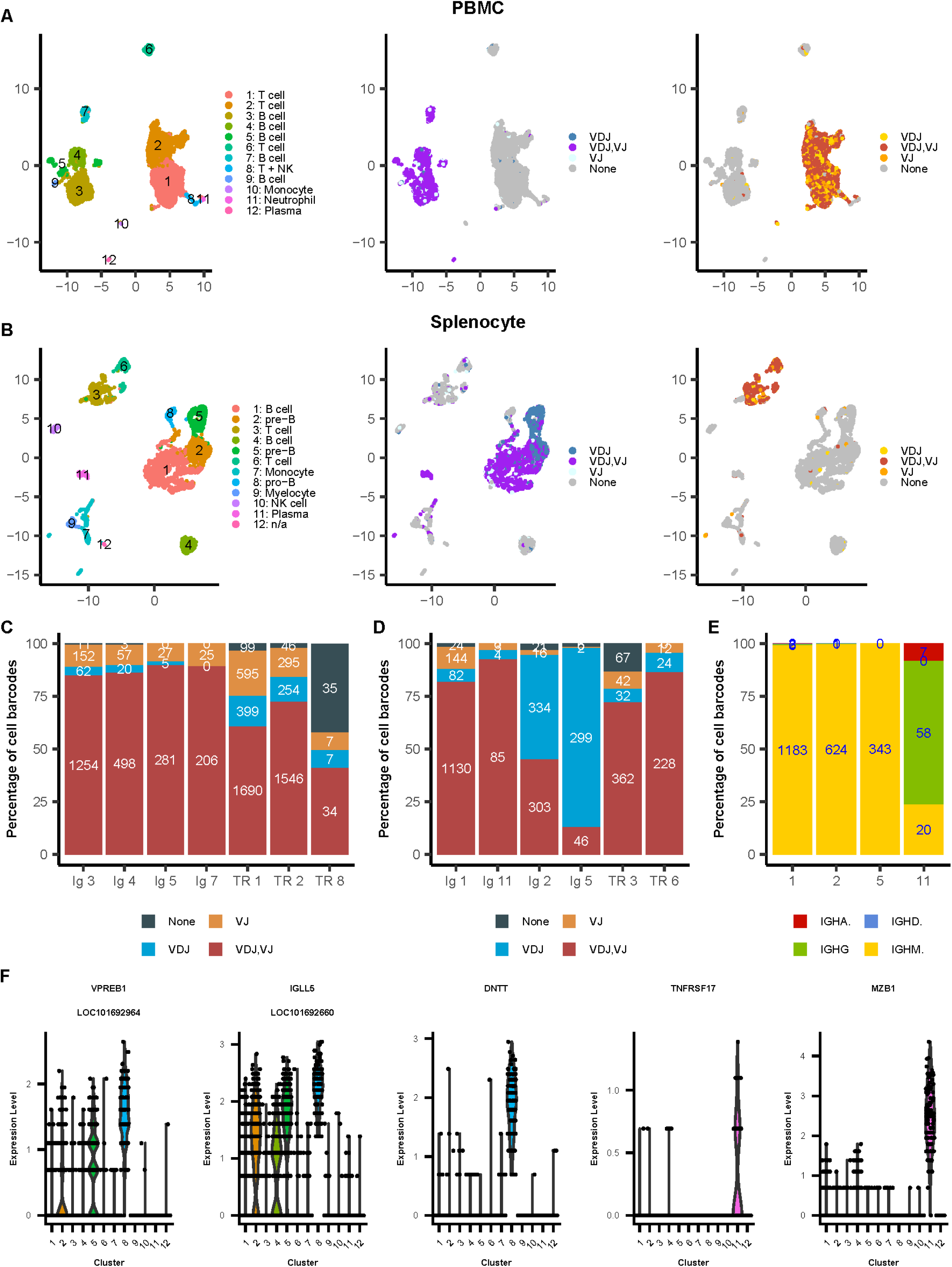
Validation of ferret single-cell gene expression and immune repertoire profiling assay. (A) UMAP plots of the (left to right) assignment of cell cluster, Ig chains, TCR chains in the ferret PBMC sample. (B) Same as (A) but for the splenocyte sample. C) Ig and TCR pairing efficiencies of the cell clusters in the PBMC sample. Percentages and numbers indicate the frequency of B and T cells with paired chains (VDJ,VJ), VDJ-only, VJ-only, or no recovered chains. D) Same as (C) for the splenocyte sample. E) IGH isotype usage of B cell clusters in the splenocyte sample shown in (D). F) Expressions of selected B cell development related marker genes in the individual cell clusters of the splenocyte sample. The cell type of cell clusters in PBMC (A) and splenocyte (B) samples was collectively annotated based on the detection of Ig and TCR transcripts, SingleR based cell type prediction, and manual review of selected canonical cell markers. The list of canonical markers used: common T cell marker: CD3D, CD3D, CD3G; common B cell marker: CD79A, CD79B, CD19; neutrophil in PBMC: S100A8, S100A9; monocyte: S100A4, LOC101672794 (PYCARD), LOC101687145 (FCN1); plasma cell: JCHAIN, TNFRSF17, MZB1; pre-B cell: surrogate light chains VPREB1 (LOC101692964) and IGLL5 (LOC101692660); pro-B cell: EBF1, DNTT; myelocyte/ immature neutrophil: MMP9, MMP8, NCF1, NCF2, NCF4; and NK cell: NCR3, NCR1 (LOC101670751). n/a in (B) indicates the cell type was not annotated due to the total number of cells (20) in cluster #12 was extremely small. For additional details, see also Figure S5 for the cell type prediction and Table S4 for differentially expressed genes in each cell cluster.

Similarly, there were 4 clusters of B cells (Figure 4B) in the splenocyte sample. The Ig pairing efficiency was 81.9% for cluster 1 and 92.4% for cluster 11 (Figure 4D). Also, most of the B cells in cluster 11 had undergone class switch, based on the high percentage of cells with IGHG and IGHA contigs (Figure 4E). The high expression of plasma marker genes such as TNFRSF17 and MZB1 also suggests that cluster 11 were likely to be plasma cells (Figure 4F), therefore with higher Ig expressions and higher Ig pairing efficiency observed. However, the Ig pairing efficiencies were much lower for clusters 2 (45%) and 5 (13%) (Figure 4D). These two clusters also included much higher percentages of cells with only IGH contigs (cluster 2: 49.6%; cluster 5: 85.7%) and had high expression of surrogate light chains VPREB1 and IGLL5 (Figure 4F), indicating that those cells were in the pre-B cell phase of development. At this stage, IGH germline sequence has undergone VDJ rearrangement, and a pre-B receptor is expressed on the surface of the cell. The pre-B receptor utilizes a surrogate light chain because neither the IGK nor IGL loci has been rearranged at this stage, and the cells express IGHM ^10^.

Cell type prediction also suggested that the cluster 8 in the splenocyte sample was a cluster of pro-B cells (Figure S5 D and E) which was consistent with the high expression of related markers such as DNTT (Figure 4F, and ^30^). This suggests that a very low detection of Ig contigs in cluster 8 (Figure 4B) was likely due to the lack of the VDJ rearrangements at the pro-B cell stage. In addition, cluster 4 in the splenocyte sample was predicted to be a cluster of B cells (Figure S5 D and E), but with a very low detection of Ig or TCR contigs as well (Figure 4B). We then approximated if the cells in cluster 4 tended to express Ig transcripts using their RNA-seq data (see Methods). We found the percentage of cells in cluster 4 that had RNA-seq reads aligned to genomic regions annotated with Ig variable genes was comparable to the B cell cluster 1 and plasma cell cluster 11, and similarly much higher than non-B cell clusters 3, 6, 7, 9, 10 (Table S5), indicating the cells in cluster 4 were very likely B cells. Interestingly, this analysis showed the pre-B cell clusters 2 and 5 had higher percentages of cells with IGH only expressions (Table S5), similarly as the VDJ analysis described above (Figure 4D). This analysis also showed the pro-B cluster 8 had a slightly higher percentage of cells with IGH only expressions, suggesting partially IGH rearrangements/expressions in those cells might have occurred. We also compared the cell gene expression profiles of cluster 4 to that of clusters 1 (high Ig pairing efficacy representing mature B cells) and 2 (high percentage of IGH only cells representing B cells in early development). Functional analysis of both lists of the differentially expressed genes showed that cluster 4 was highly enriched with cell death related biological functions (Table S6). Since we were not able to assemble full-length Ig transcripts through V(D)J sequencing, these results suggest that the overall quality of RNAs including Ig transcripts of the cells in cluster 4 could be too low, either due to low cell quality or cell death triggered by incomplete VDJ/VJ rearrangements. Cluster 10 in the splenocyte sample was predicted to be a cluster of T and NK cells (Figure S5 D and E), but the high expressions of NK cell markers such as NCR3 and NCR1 (LOC101670751) and low expressions of T cell markers such as CD3E and CD3G suggest that cluster 10 is most likely to be predominately NK cells (Figure S5 F).

Thus, the observed variations in ferret Ig and TCR pairing efficiencies corresponded to the expected biological functions and developmental progression of cells in these compartments, demonstrating that our analyses are able to elucidate the Ig and TCR transcript abundances in individual ferret B and T cells and link them to their developmental stages. The resources developed here are therefore suitable for efficient single-cell based immune repertoire sequencing analysis of ferret cells.

## DISCUSSION

Ferrets are a valuable small animal model for several human diseases, including influenza, SARS-CoV-2, and cystic fibrosis. However, research with this model has been limited by a lack of reagents and genetic information, especially as immune repertoire studies have become widely used in the studies of humans and other species. In this study, we have identified and annotated both constant and variable regions for ferret Igs and TCRs. We used long read transcriptome sequencing, combined with available genome sequences, to generate the first complete reference of C-regions from all expected ferret Ig and TCR isotypes and chain types, as well as an extensive reference annotation of V-, D-, and J germline genes. We also developed and experimentally validated the first ferret-specific single-cell paired gene expression and immune repertoire profiling assays compatible with the high-throughput 10x Genomics platform. These results mark a major advancement in ferret immunogenetic resources and potential for improved translational medicine regarding domestic ferrets as an animal model of infectious diseases.

We find two TRBC genes in ferrets, as did Gerritsen et al. ^21^, which is also the known number of TRBC genes in human, dog, rhesus macaque, and cat ^31^. We also identified ten TRGC genes, of which six are functional; ^31^dogs have six functional TRGC ^32^. IGHG is also known to have multiple subclasses, including four in dogs ^33^, four in mink ^34^, and three in cats ^35^; we find two IGHG subclasses in ferrets. The genomic version of one of these subclasses (IGHG2) is classified here as a pseudogene (Table S2) due to apparent frameshift mutations in the sequence; however, these may represent sequencing errors, since mRNA transcripts of this gene with frameshifts corrected (here referred to as IGHG2*02 and IGHG2*03) are present at high levels in transcriptomes, while IGHG2*01 (the frameshifted version) represents only 0.15% of the total transcripts of secreted IGHG2 (Table 1, Table S2 “Allelic Variants” tab). No exact matches for the genomic sequences of IGA were found in transcriptomes; this may reflect an allelic variant not present in the ten ferrets analyzed, or may also reflect a sequencing error in the genomic sequence. In humans, there are seven IGL genes, of which three (IGLC 4, 5 and 6) are pseudogenes ^36^. Ferrets also have seven IGLC genes, positioned in the same arrangement as in the human genome (Figure 1C); however, only two (IGLC 4 and 6) seem to be pseudogenes, since unlike the human situation, no premature stop codon is present in IGLC5 and the splice signal is intact. The sequences of IGLC3 and IGLC5 are identical, and the sequences of the proteins encoded by IGLC1, IGLC3, and IGLC5, are identical (Table S2).

We identified both membrane-bound and secreted forms of all IGHC, differentiated by distinct exon usage at the 3’ end of transcripts. As expected, IGA and IGG transcriptomes were dominated by secreted versions of the genes, while transmembrane versions of IGM and IGD were more commonly seen in transcriptomes. Interestingly, an allelic variant of IGD (IGHD*02) was identified in transcriptomes containing an 18-base pair insertion in the transmembrane exon (Table S2 “Allelic Variants” tab). Unsurprisingly, IGE expression in splenocytes from these young, healthy, naïve, laboratory-housed ferrets was very low. Similarly, TCR expression in transcriptomes was generally much lower than for immunoglobulins, with some subclasses being undetected; allelic variants of TCR constant genes very likely are present in ferrets as well, but the low counts make them difficult to confidently identify. Supporting this, a previously published TRBC sequence (here termed “TRBC1*01”) differs by one base pair from the sequence identified here (“TRBC1*02”; Table S2 “Allelic Variants” tab), and presumably these are allelic variants at this locus. Additional alternative splicing events that were identified were at low frequency in the lymph node and splenocyte transcriptomes and included retained introns, skipped exons, and variation in the start positions of exons, including some apparent scFv sequences. The abundances in other tissues and the biological function, if any, of these apparent splice variants requires more studies.

For each of the constant regions (IGHC, IGKC, IGLC, TRAC, TRBC, TRDC, TRGC) we identify V and J genes (and where expected, D genes) in the expected genomic contexts (Figures 1-2; Figure S1). Genome analysis also identified variable genes in the TRB and TRD region without corresponding constant regions (Figure 2B,C; Figures S1). In some cases, putative VH genes were identified on a contig that did not overlap with constant regions, so that their genomic context could not be accurately determined. These genes were assigned provisional names pending more genome data, and are listed in Table S2, “IGHV Provisional” tab.

The degree of overlap between the VDJ-region genes we identified here and those reported by others varied among the different IG and TR regions. For example, Wong et al^20^ identified 29 IGHV, 53 IGKV, and 34 IGLV genes, while this study identifies 82 (including 19 provisional genes: Table S2, “IGHV Provisional” tab), 87, and 136 respectively, including genomic context and additional sequence information (e.g., leader sequences) that was not available to previous work. Gerritsen et al^21^ identified 27 TRBV, 2 TRBD, and 12 TRBJ genes, and interestingly we did not identify any new genes in the TRB region. In some cases, putative Ig or TCR V-region genes that did not meet our filtering criteria did overlap with the sequences reported by Wong et al^20^, or Gerritsen et al^21^ and these may represent allelic variants (given the outbred nature of ferrets used in these studies) or pseudogenes. In cases where genes identified here are identical to those previously identified (e.g. 21 of the 27 TRBV genes described by Gerritsen et al^21^), we have noted this in a “Comment” column (Table S2, “Previously Identified” tab); it is likely that some non-identical genes also represent allelic variants of previously described genes.

Although we examined transcriptome data only from a limited number of ferrets from a single supplier, allelic variants in the C regions were readily detected in these datasets (e.g. 4 alleles of IGHA, 3 of IGHE and IGHG2, and 2 each of IGHD, IGHG4, and IGHM: Table 1; Table S2, “Allelic Variants” tab), including both point variants (e.g. IGHA1-1 vs IGHA1-2) and insertion/deletion variations (e.g. a 6 amino acid insertion near the C-terminus of the membrane-anchored form of IGHD1*02 vs IGHD1*01). In addition, several sequences identified here are ∼99% identical to those identified previously and in the same genomic context ^20, 21^, and are therefore potentially allelic variants of those reported previously. It is likely that much more allelic variation exists in the worldwide population of laboratory and domestic ferrets. We did not attempt to identify V/D/J allelic variants in this study, because factors such as hypermutations may complicate the accurate detection of allelic variants in the rearranged Ig/TCR transcript sequences. However, it is likely that V/D/J allelic variants are also common. Consensus sequence generation of the most abundantly expressed IGHV, IGKV, and IGLV genes revealed some variation between alleles and subclasses, including in the positions of the primers designed by Wong et al ^20^. This allowed us to consider sequence variation in C-region genes when we designed our ferret-specific assays, targeting positions within Ig and TCR C-region genes that were characterized by no or minimal sequence variation across different subtypes, isoforms, and alleles. We demonstrate that these primers amplify a substantial number of highly abundant V-region genes, as well as multiple lesser-abundant V-region genes, which are inefficiently amplified by previous Ig MPCR assays. Although the primers designed by Gerritsen et al ^21^ were able to achieve 100% amplification efficiency for each TRBC sequence in our reference, the position and melting temperature parameters of these primers were not compatible with the 10x Genomics immune profiling assay. We were able to target and enrich every Ig and TCR chain and isotype and recover B and T cells with matching transcriptomic profiles and paired repertoire profiles. Analysis of these two data together provides valuable insights into the functionality and development of B and T cells. This was exemplified by our analysis of IGH-only B cells detected in our splenocyte sample.

By combining genome analysis with full-length transcriptome sequencing data, we were able to greatly increase the number of annotated Ig and TCR V-, D- and J-region genes, including over 100 new Ig V-, D-, and J-region genes. The ability to annotate the V, D, and J genes of Ig and TCR transcripts correctly and robustly is critical to future single-cell immune repertoire profiling assays. Overall, the use of these annotations and single-cell sequencing assays provides a valuable resource to infectious disease researchers utilizing the domestic ferret animal model.

### Limitations of the study

There may be several potential limitations in this study. First, we directly sequenced the lymph node and spleen samples using Iso-seq, and consequently the percentages of sequencing data representing Ig and TCR transcripts were low. In some cases, it was challenging to accurately construct reference sequences and we had to rely on additional analysis such as transmembrane domain search in CCS reads. Using isolated B and T cells will significantly improve the coverages of Ig and TCR transcripts, and therefore reduce the computational challenges and likely improve the quality of the final references, but would increase the experimental complexity for a species lacking reagents. Also, the detection and annotation of Ig and TCR genes could be biased toward those genes expressed and detected in the specific samples used in this study. Considering that IMGT continues to curate and update its databases with newly discovered germline V, D, and J gene sequences for well-studied genomes, such as those of humans and mice, it is very likely that additional ferret V, D, and J genes will be identified. Second, the allelic variants we observed were based on the analysis of transcript sequences which may not have even representation across all positions. DNA sequencing will be more accurate and lead to more complete results. Third, we annotated V, D, J genes on an unpublished draft ferret reference genome assembly. Even though this current reference genome assembly used third generation long read sequencing technology and is significantly more continuous than the previously released ferret assembly MusPutFur1.0 (an increase of contig N50 size from 44.8 kb to 23.6 Mb), it is not a haplotype-resolved genome assembly. Therefore, some of the annotated alleles may represent potential heterozygosity or a collapse of duplicated segments. The sequences of some constant genes were not found in the analyzed cDNA datasets and this could indicate potential sequencing errors. We successfully identified 57% of V, D, J non-pseudogenes of the entire ferret germline repertoire expressed in full-length and 100% identity in cDNA sequences (Table S2). In the future, the generation of haplotype-resolved ferret genome assemblies using longer and higher quality reads with improved assembly algorithms will continuously enhance the ferret immune repertoire.. Fourth, for the single cell analysis we only analyzed PBMC and splenocyte samples from naïve animals. Additional samples from infected and/or vaccinated animals may provide additional information about ferret Ig and TCR repertoires. Fifth, we performed ferret cell type prediction and classification based on the human references and a small number of canonical human immune cell markers, but we did not perform detailed benchmark analysis of the accuracy of these predictions.

## METHODS

### Resource availability Lead contact

Further information and requests for resources and reagents should be directed to and will be fulfilled by Lead Contact, Xinxia Peng.

### Materials availability

This study did not generate new unique reagents.

### Data and code availability

All ferret Iso-seq and single-cell RNA-seq data were deposited into NCBI (BioProject accession #: PRJNA939558 and GEO accession #: GSE231948) and are publicly available as of the date of publication. All ferret C, V, D, and J sequences are available in the IMGT database under accession numbers IMGT000135, IMGT000136, IMGT000131, IMGT000203, IMGT000195, IMGT000196, IMGT000168 or NCBI TPA accessions BK063796, BK063797, BK068009, BK068010, BK068295, BK068011, BK068537 . Code is available on Github: https://github.com/ncsu-penglab/FerretIgTCR.

### Method details

### Ferret tissue sample preparation and transcriptome sequencing of Iso-Seq analysis

All ferrets originated from Triple F Farms LLC (Gillett, PA). To construct cDNA libraries for Pacific Biosciences (PacBio) single molecule real-time (SMRT) sequencing, RNAs were isolated from two tissue types: lymph node (LN) and splenocytes. Splenocyte samples were from 4-6 month old female specific pathogen free (SPF) ferrets that were serologically negative for influenza (used under CDC IACUC protocols #2885YORFERC and 3168YORFERC). Lymph node samples were archived samples from 16-19 weeks old male and female ferrets in a bartonella infection study (used under NC State University IACUC #18-175-B). Additional information of the ferrets and samples including source, age, sex, and infection/treatment is in Table S1.

Tissue from five colonic and three cranial mesenteric lymph nodes were flash frozen in liquid nitrogen. No more than 100mg of frozen tissue was placed in at least 10 volumes of RNAlater ICE (Invitrogen) overnight at -20C. After overnight incubation, tissue samples were transferred to a 2mL tube containing 2.8mm ceramic beads and 1mL TRIzol (Invitrogen) for RNA isolation. Samples were processed in a FastPrep 24 (MP Biomedicals) for 2-4 cycles of 20s at a speed of 4.5 with 1 min ice rests between cycles. Exact cycle numbers were determined based on visual observance of tissue homogenization. Homogenized samples were isolated according to the TRIzol procedure for RNA to the point of isolating the aqueous phase, after which each phase was precipitated with 1.5X volumes 100% ethanol. Samples were then applied to RNeasy (Qiagen) columns, treated with DNase, and isolated according to the manufacturer’s protocol. Samples were quantified using a NanoDrop (Thermo) and were assayed for RIN using a Bioanalyzer (Agilent). Only samples with a RIN ≥7.5 were used for Iso-Seq library construction. Iso-Seq library construction was performed using 300ng input RNA per sample following the manufacturer’s specification (Iso-Seq Express Template Preparation for Sequel and Sequel II Systems, Iso-Seq Express 2.0 Workflow) and was sequenced using a Sequel II System (Delaware Biotechnology Institute at the University of Delaware). RNA samples were barcoded separately and pooled together for sequencing.

Ferret splenocytes were stored in liquid nitrogen in the presence of 10% DMSO and then processed in a similar manner to the lymph nodes. Total RNA from splenocytes was extracted with the RNeasy Micro Kit (QIAGEN). High quality RNA (with an RNA integrity number over 7.0) was processed for cDNA synthesis by following the Iso-Seq express template preparation protocol for Sequel and Sequel II sequencing systems. cDNAs were amplified and those with a size greater than 2kb were selected with ProNex beads (Promega). cDNA SMRTbell libraries were prepared and sequenced with a PacBio Sequel II system in the core facility at Centers for Disease Control and Prevention. RNA samples were barcoded separately and pooled together for sequencing.

Raw PacBio Iso-Seq sequencing data were first run through the circular consensus sequence (CCS) protocol in SMRT Link v8.0 to generate CCS reads. Each CCS read represents the consensus sequence from multiple passes of a single transcript. CCS reads were classified as non–full length and full length, in which the latter has all the following detected: polyadenylation tail, 5’ cDNA primer, and 3’ cDNA primer.

### Identification and annotation of ferret Ig and TCR constant regions

Here we aimed to generate the complete cDNA sequences representing the entire ferret C-region (i.e., including the 3’ UTR) from the Iso-Seq data. We first identified putative full-length ferret Ig and TCR transcript sequences by aligning all full-length ferret Iso-Seq CCS reads to a custom Ig/TCR database using IgBLAST (v1.16.0) ^37^, with an e-value cutoff of 0.01. Due to the incompleteness of the domestic ferret reference sequences in the IMGT database, before the finalization of this project, our custom IgBLAST database was composed of Ig and TCR V, D, J sequences from several related species including dog, cat, horse, and TRB. Full-length CCS reads were retained as putative Ig/TCR sequences for downstream analyses if they had a variable gene hit with an e-value < -log_10_(50) and a joining gene hit < - log_10_(3).

Next, we extracted the C-region sequence from each of the putative full-length ferret Ig/TCR transcript sequences, based on the J-region end position reported by IgBLAST alignment. Extracted C-region sequences of high similarity were clustered together using CD-HIT v4.6.6 ^38^ with the following parameters: -c 0.99 -G 0 -aL 0.95 -AL 100 -aS 0.99 -AS 30 -d 0 -T. We only kept the clusters with at least 10 extracted C-region sequences for the consensus sequence generation. After clustering, a consensus C-region sequence was generated from each cluster. Abundant allelic variants, i.e. alternative alleles represented by approximately 50% of CCS reads that were uniquely aligned to the corresponding position on the reference genome assembly with more than 98% identity, were interpreted as allelic variants. We used the cons tool from the EMBOSS v6.6.0 suite ^39^ to generate consensus sequences, identifying and removing any remaining insertions at predominantly gapped locations. To ensure that these consensus sequences were representative of true C-region sequences captured by our Iso-seq analysis, we assessed the quality of the generated consensus sequences by calculating the percentage of consensus at each base position and insertion and deletion rates. We removed all consensus sequences from our analysis if they had more than 10 base positions with a consensus percentage less 90%, an insertion rate above 0.0015 or a deletion rate above 0.0007.

To remove potentially redundant C-region consensus sequences, we performed a second clustering and consensus sequence generation step after aligning them to the recently released ferret reference assembly (NCBI accession no: GCA_011764305.2, generated by JCVI from the haploid genome of an outbred adult male ferret, using a combination of short and long reads ^24^). The corresponding NCBI RefSeq genome assembly was GCF_011764305 ^23^. This recent ferret genome assembly appears to be more continuous (contig N50 of ∼24M bps vs. ∼45k bps) than the previous published draft ferret genome sequences ^40^. We aligned the ferret C-region consensus sequences generated above to the reference genome using the minimap2 alignment tool ^41^ with parameters: -ax splice:hq -uf --secondary=no. This round of ferret C-region consensus generation was performed with the collapse_isoforms_by_sam.py script from the Iso-seq bioinformatics platform (https://github.com/Magdoll/cDNA_Cupcake) to produce full-length, non-redundant consensus sequences. After this round of consensus sequence generation, we remapped these updated ferret C-region consensus sequences to the same reference assembly similarly using minimap2, and visualized the alignments in the Integrative Genomics Viewer (IGV) browser ^42^. Aligned ferret C-region consensus sequences were manually curated based on intersecting alignments of annotated C-region sequences from IMGT.

To check that these Iso-Seq data derived ferret C-region transcript consensus sequences are functional we performed two assessments. First, we generated amino acid sequences by translating the C-region consensus sequences using the EMBOSS transeq tool ^39^, and kept the frame with the longest amino acid sequence for each consensus sequence. Amino acid sequences were then analyzed using the Pfam collection of protein families ^43^ for functional domain prediction, particularly the presence of C1-set or DUF1968 protein domains. C1-set domains are found in immune system molecules such as Igs and TCRs. DUF1968 domains are representative of the TRA C-region domain. Second, we manually examined the corresponding donor and acceptor splice site sequences, after aligning these C-region consensus sequences to the ferret reference genome assembly. Since we observed both a high rate of predicted C-region domains (C1-set & DUF1968) in the translated amino acid sequences and corresponding canonical donor (GT) and acceptor (AG) splice sites, we removed those consensus sequences without a predicted C1-set or DUF1968 domain from further consideration.

Immunoglobulins can be secreted or membrane bound. Therefore, we further classified our IGH C-region consensus sequences as membrane bound and secreted isoforms for each IGH isotype, based on the presence or absence of a transmembrane domain. To do this we generated translated amino acid sequences from each consensus sequence using transeq ^39^ and inputted them to the TransMembrane Hidden Markov Model (TMHMM) (v2.0c) sequence analysis tool ^44^ or DeepTMHMM ^45^. To thoroughly identify corresponding membrane-bound isoforms we first aligned all CCS reads to their respective secreted isoforms (see below) to identify the corresponding gene. We then scanned the aligned CCS reads for the presence of transmembrane domains similarly. This analysis also allowed us to identify a membrane bound IGHE isoform. This additional sequence was similarly evaluated as described above and added to our C-region reference set.

We also assessed the completeness of this new ferret C-region reference set by comparing the overall consistency in identifying putative Ig and TCR transcripts in the Iso-Seq data using the custom IgBLAST with variable region reference sequences from related species vs. aligning directly to this new ferret C-region reference databases. We independently classified all CCS reads as putative Ig and TCR transcripts as follows. Iso-Seq CCS reads were globally aligned to this new collection of ferret Ig and TCR C-region sequences using the USEARCH - usearch_global tool ^46^ with a 90% identity threshold to ascertain chain type and isotype/subclass where appropriate. CCS read sequences without large gaps (>10 bp) in their alignment were kept in the final assignment of Ig and TCR transcript sequences, as it was unclear if those sequences with larger gaps were rare alternatively spliced transcripts or the result of sequencing errors. The agreement was evaluated by the overlap between the CCS reads captured by the new ferret C-region reference databases with those identified by the initial IgBLAST search.

The final set of C-region reference annotation was further reviewed by extensive manual curations including the addition of two IgL pseudogenes and named by the corresponding human nomenclature.

To assess the sequence quality of putative Ig/TCR transcript sequences initially identified by IgBLAST, we extracted the C region portions from the Ig and TCR CCS reads based on IgBLAST and aligned the extracted C region portions against the final set of C-region reference sequences using the USEARCH usearch_global tool with parameters: -id 0.65 -strand both - gapopen 10.0I/0.0E -gapext 1.0I/0.0E --query_cov 0.99. We then quantified the mismatch, insertion, and deletion events within these alignments.

### Initial annotation of ferret Ig and TCR germline V and J genes

Similarly, we started annotating germline V and J genes by leveraging the full-length Ig and TCR transcripts in the ferret Iso-seq data. V-region sequences were extracted from Iso-Seq CCS reads using the start and end position reported by the custom IgBLAST described above. We considered the entire sequence up to the V-region end position. Therefore, our putative V-region sequences contained the 5’ UTR, first leader sequence (L1), second leader sequence (L2), and V-region sequences (FR1 to CDR3). To determine the genomic positions of germline V and J genes, we aligned the putative V- and J-region sequences from CCS reads to the ferret reference assembly (NCBI accession no: GCA_011764305.2) using the minimap2 sequence alignment tool (56) with parameter-ax splice:hq -uf --secondary=no. At this step we did not allow for secondary alignments as we reasoned that the germline sequences located by the primary alignments would have the best chance of locating authentic V genes.

Given the expected genomic organization of V-region genes, we filtered the alignment data to only include sequences with mappings consistent with the genomic structure of these genes. V genes are composed of two exons. The first exon is the L1 sequence, and the second exon is composed of the L2 and V-region sequences ^47^. We observed that more than 80% of our putative V-region sequence alignments had two exons and therefore removed the other 20% with more or less than two aligned exons. Additionally, we observed a small number of V-region sequence alignments that had an unusually large intron length (>350 bps) and a large first exon length (>200 bps). These alignments were also removed from the downstream analysis. The values of these criteria were chosen empirically, and sequence alignments outside of these values represented rare outliers. When multiple extracted V-region sequences were mapped to the same genomic locations, overlapping sequence alignments were collapsed into single loci using the merge command from bedtools ^48^.

Further, we considered the potential of these putative V genes producing functional V(D)J transcripts and receptor proteins by the presence of a downstream RSS and a complete ORF. We examined the 50 bps directly downstream of the 3’ end of the putative V genes for the presence of an RSS. These 50 bp sequences were scanned for the presence of an RSS using the RSSsite software ^49^. To mitigate potential false negatives in the analysis of putative RSSs, we also considered whether the putative V gene had a complete ORF. To do so, we performed a three-frame translation of putative V gene sequences using the EMBOSS transeq tool. Because the captured L1 sequence represented the beginning of the V(D)J coding region, we required that each ORF have a methionine amino acid and removed any amino acid sequence upstream the first methionine. Next, we removed any sequence downstream of the first stop codon (if one was detected) in each translation frame. We selected the translation frame with the longest ORF for each putative V gene. Based on these two criteria, we removed sequences that did not have a downstream RSS detected and did not have a complete ORF, i.e. free of stop codons from the methionine in L1 to the detected RSS.

Annotation of J genes was performed similarly to the V genes. We extracted J-region sequences from Iso-seq CCS reads similarly using the method described for V- and C-region sequences. However, we limited this analysis to only CCS reads that were both reported by the custom IgBLAST and mapped to our ferret V-region and C-region references. The alignment of the extracted putative J-region transcript sequences to the ferret reference assembly was done using GSNAP (2017-04-13) ^50^, a mapping tool for short read sequence alignment. The ferret reference assembly sequence was also filtered to contain only regions in between and including V- and C-region genes. This narrowed the search of germline J genes to genomic regions flanked by annotated V- and C-region genes. Due to the short sequence length, our J-region filtering criteria was relatively less stringent. We required that there be only one exon per J-region sequence alignment. This criterion was consistent with over 99.9% of our J-region sequence alignments. Overlapping alignments were similarly merged and filtered based on the presence of an upstream RSS. Additionally, we observed that in some cases the upstream RSS was within the putative J-region gene body, so the range of the RSS search consisted of 50 bps upstream of the J gene start and up to 50 bps within the J gene body.

### Annotation of additional ferret Ig and TCR germline V and J genes

One potential constraint of the above transcript alignment based analysis is that only a portion of the ferret Ig and TCR germline genes may be annotated, since not all V, D, and J genes were expressed and therefore captured by our Iso-seq data. In addition, the presence of extensive somatic mutations in immunoglobulin variable regions also means these regions may not be located directly from transcript alignments. Therefore, we identified and annotated additional germline V and J genes as follows.

We first iteratively scanned the reference genome assembly for additional Ig and TCR V and J genes that may have been undetected initially. For each iteration we utilized the current set of annotated V and J genes as references, as we expected high sequence similarities among genes in the V(D)J region in general. To search for additional V and J genes in the same genomic region, we first masked out the currently annotated loci using the bedtools command maskfasta. Next, we aligned the current V- and J-region genes to the reference genome assembly using the same procedures described initially, but with less stringent scoring parameters. All new alignments were analyzed and filtered using the same RSS and ORF criteria described above. The only difference to our filtering procedure was that we used a custom R script to score and classify putative RSSs rather than RSSsite ^49^. The presence of an RSS was determined using the same formulas and thresholds described in ^51^. We confirmed the appropriateness of the score thresholds based on the assessment of the RSS score distributions of randomly selected sequences in the ferret genome that started with a CA, similarly in previous studies ^49, 51^. Based on the RSS and ORF criteria we classified new sequence alignments as true or false positives. True positives were added to our reference set, but both true and false positives were masked in the next iteration. We repeated this masking, searching, and filtering procedure until no new true positives were detected.

In parallel, for additional Ig germline V genes we used the VgenExtractor tool ^52^ to locate potential V-exon regions on the ferret genome assembly. Using a Python script, the genomic region 500 bps upstream of each putative V gene was checked for the presence of probable leader sequences, and the region 55 bps downstream was checked for probable RSS sequences. The extended genomic sequences obtained were translated in all six frames using EMBOSS sixpack software ^39^ and manually examined. The upstream genomic regions corresponding to the Iso-Seq Ig V leader sequences were annotated as the L1 region.

To identify additional Ig J genes, we generated a list of non-redundant reference sequences for all immunoglobulin heavy and light chain J-region protein sequences in IMGT/GENE-DB ^53^. A local BLAST ^54^ database was created using the contig sequences that contained most of the IGH, IGK and IGL V genes along with the C region genes from the ferret reference genome assembly, since those contigs were expected to also contain the corresponding germline J and D genes. A translated BLAST search was performed for each of the IMGT J genes against this local BLAST database. The genomic regions with a minimum match percentage of 60% were retained as putative J genes and the 50bps upstream and downstream of these putative regions were checked for the presence of RSS sequences using the Python script.

### Annotation of ferret Ig and TCR germline D genes

Since the D genes are extremely short therefore challenging to align directly, we did not utilize the IgBLAST alignment of Iso-seq CCS reads to annotate D-region sequences. Instead, we devised two distinct but complementary approaches to annotate ferret germline D genes. For the first approach, we utilized a similar iterative RSS based scanning strategy as described above to delineate the start and end positions of D genes, based on their expected gene structure. In brief, TRBD and TRDD genes are flanked on the 5’ end (V-D) by an RSS with a 12 bp spacer on the opposite strand as the D gene and an RSS with a 23 bp spacer on the same strand at the 3’ end (D-J). IGHD sequences are flanked by 12 bp spacers on both the 5’ and 3’ ends in the same strand orientations as TRBD and TRDD. We therefore utilized the 5’-RSS-D-RSS-3’ organization to determine the D gene start and end positions at the IGH, TRB, and TRD loci.

Scoring of putative RSS sequences was done using the same procedure employed in the iterative genomic search of V- and J- region genes as described above. If two adjacent RSS sequences were less than 50 bps apart and in the expected orientation, then the sequence in-between the two RSSs was classified as a D gene.

For the second approach, Ig D regions were first estimated within the VDJ-rearranged Iso-Seq CCS reads by searching for an extended CDR3 sequence pattern “Y[HY]C.{1,55}VSS” that incorporates the flanking residues, in addition to the previously reported CDR3 region ^55^. The estimated D-region protein sequences were queried against the same local BLAST database as described above containing D genes using translated BLAST. The genomic regions with a minimum match percentage of 60% were retained for further analysis. The length of germline D genes in IMGT ^53^ varies from ∼3 – 15 amino acids and chances for picking random matches in the shortlisted blast hits are expected to be high. Accordingly, to reduce false positives in D gene selection, combinations of 3bp patterns based on RSS conserved nucleotides like ‘CAC’ from heptamer and ‘ACC’ or ‘AAA’ from nonamer were selected and searched on both the sides of the putative D genes. The results were further evaluated manually to exclude D genes with defective RSS flanking sequences. ^18, 19^

Finally, amplicon sequence sets were directly submitted to IMGT/HighV-QUEST ^56, 57^ which implements IMGT/V-QUEST program version 3.6.3 (January 2024, 30) and IMGT/V-QUEST reference directory release 202430-2. To focus on Ig/TCR transcript sequences, transcriptomic reads were first assembled using MiCXR ^58^ v4.6.0 using a customized library created from IMGT reference directory 202430-2 (23 July 2024) for ferret species before the analysis by IMGT/HighV-QUEST. The IMGT/HighV-QUEST results are provided in 11/12 CSV files the content of which have been previously published ^59^. Productive rearrangements were studied to analyze the expression of V and J genes with an ORF functionality due to noncanonical V-RS sequences. For the analysis of the expression of ORF D-GENES, the CIGAR format for the D-REGION generated by IMGT/HighV-QUEST in vquest_airr.tsv file was used to identify non-trimmed and non-mutated rearranged D genes.

### Classification of germline V, D, J genes as functional or pseudogenes

Pseudogenes for V-gene units are commonly found in genome sequences, but do not contribute to functional Ig or TCR. We identified pseudogenes based on IMGT criteria ^47^. Briefly, a V gene was considered functional if it (i) lacked a stop codon or a frame-shift mutation in the leader sequence part1 and V exon, (ii) contained the conserved tryptophan and cysteines, (iii) contained no defect in the splicing sites, recombination signals and /or regulatory elements, and (iv) had an intron less than 500bps between the leader sequence and V exon. V genes that met these criteria but that lacked any of the three conserved amino acids were classified as “ORF”; those that failed to meet the criteria were classified as pseudogenes.

### Nomenclature of V, D, and J genes

Genes were assigned names as previously described ^15, 28, 58^. Briefly, V genes were subgrouped by comparison to human sequences and by 75% percent identity at the nucleotide level. They were assigned a number for the subgroup, followed by a hyphen and a number for their localization from 3′ to 5′ in the locus. The J and D genes were similarly assigned to a subgroup according to the human sets, followed by a number based on genome location. Constant regions were assigned names according to homology to human sequences, while homology to canine sequences was used to resolve ferret IgG2 and IgG4 constant regions.

### Design of ferret-specific primers targeting Ig and TCR C-regions

When designing ferret-specific primers to target and enrich Ig and TCR mRNA transcripts we wanted to account for all observed variations in these sequences, including allelic, isoform, and subtype. To account for variation between different animals (alleles) in C-region genes, we used all 10 additional domestic ferret genome assemblies available under the same NCBI BioProject (accession: PRJNA580247) as the ferret reference genome assembly. Of these 11 publicly available genome assemblies 10 were sequenced using Oxford Nanopore PromethION technology and one was sequenced using both Illumina NovoSeq 6000 and PacBio RS technology. Additionally, two of the 11 genome assemblies were generated using DNA samples from Marshall Farms’ animals and the other nine were taken from Triple F farms.

We aligned the annotated C-region reference sequences as described above to each of the additional 10 genome assemblies using the minimap2 ^41^ tool with parameters: -ax splice:hq -uf – secondary=no -G20k. The resulting alignment files were converted to bed12 files using the bedtools (v2.30.0) ^48^ command bamToBed with parameter -split to account for splicing in C- region genes. Bed files were then used to extract the genomic C-region sequence for each gene from each assembly using the bedtools getfasta command with parameters -split -s to account for the strandedness and splicing of the sequence.

For each C-region gene a consensus sequence was generated with the protocol described above with 11 references, one from each genome assembly. Rather than designing separate primers for each subtype and isoform of a C-region gene, all the C-region sequences of the same chain type (or isotype for IGH) were used to generate a consensus sequence. Only the first 500 bps of each sequence were used in consensus sequence generation and considered in the primer design for the 10x Genomics single-cell immune profiling assays. Next, we remapped the 11+ (>11 if more than one isoform or subtype) reference sequences to our consensus sequences using USEARCH -usearch_global to measure the percentage of consensus at each position. To ensure that our primers would be able to enrich for all alleles, consensus sequences were altered such that every base pair with less than 100% consensus was masked (ACTG → N). In this case, the primer design software Primer3 (64) would only consider sections of consensus sequences that were identical across all 11+ reference sequences. We additionally restricted our primers to having a melting temperature between 60 and 70 degrees and a GC content between 50 and 60%. In some cases, primers necessitated manual design to achieve desired parameters and capture.

After primer design and selection, we again tested the ability of our primers to amplify each of the 11 C-region references using in-silico PCR (isPCR). Across every Ig and TCR gene in our reference we were able to amplify all Ig and TCR C-region genes identified on the 11 assemblies in silico with a 100% primer match with the exception of TRGC1*01, TRGC2*01, TRGC6*01, and TRGC7*01. TRGC6*01 is a pseudogene with a truncated exon 1 sequence that does not have the inner primer binding site. Each of TRGC1*01, TRGC2*01, and TRGC7*01 contains one mismatched base at position 5 or 6 for the inner enrichment primer. This mismatch was necessary to achieve a complete complementary pairing with other TRG constant regions. Given the placement of this mismatch near the 5’ end of the primer and the presence of the mismatched primer in the second enrichment reaction, the recovery of these three TRGs would still be possible with this design.

### Validation of ferret-specific primers targeting Ig and TCR C-regions

We first evaluated ferret-specific MPCR assays for amplifying IGH, IGK, and IGL transcript sequences in the Iso-Seq data *in silico*. The input sequences to these assays were the ferret Iso-seq CCS reads used throughout this study that met filtering criteria after the custom IgBLAST alignment. We considered a transcript to be amplified after only a single round of amplification despite the nested RT-PCR design of this approach. The Wong et al. ^20^, Gerritsen et al. ^21^, and our newly designed reverse primers used were also assessed via *in silico* PCR analysis in which a dummy adapter sequence was prepended to the 5’ end of sequences to enable testing the differences in the reverse primer efficiency using the University of California, Santa Cruz isPcr tool ^60^. These *in-silico*, template-switch PCR experiments were performed on the 11 ferret genome assemblies for each C-region gene. For the isPcr tool we used the parameters which required a 14-nucleotide perfect match from the 3’ end of the primer (-tileSize=11 -minGood=14 -minPerfect=14). The adjustment from the default value of 15 to 14 nucleotide perfect match from the 3’ end of the primer was made to amplify the pseudogene TRGC2*01 *in silico*.

Ferret Ig and TCR specific primers were further experimentally validated against cDNAs isolated from a pool of three PBMC samples, each from a different animal. For each ferret Ig and TCR specific primer, a corresponding forward primer was designed within the region upstream of the C-region reverse primer. These forward primers were then paired with the appropriate assay reverse primers and assessed for specificity via PCR in separate reactions and in pools representative of their final concentrations in the 10x Genomics VDJ enrichment protocol. PCRs were performed at an annealing temp of 65°C using Accuprime Taq polymerase (Invitrogen) with 1 minute extension and 30 cycles. All primers were additionally tested with qPCR using PowerUp SYBR Green (Applied Biosystems) to confirm expression levels and rule out the possibility of significant primer bias occurring in the pooled reactions. PCR products of such reactions were analyzed on a 1% agarose gel pre-strained with SYBR-Safe to confirm the amplification specificity.

### Ferret tissue processing and cDNA generation for single cell sequencing

For single cell sequencing analysis one cryopreserved PBMC and one splenocyte sample were randomly selected from two influenza-naïve male animals between 1.3-1.5 years of age (CDC IACUC protocol #3139ROWFERC). Cells were washed in resuspension buffer (RPMI, 10% FBS), depleted of dead cells using a removal kit (Miltenyi Biotec), collected by centrifugation at 700xg, and resuspended at an appropriate concentration, between 700 and 1200 cells/µl. All cells were counted by hemocytometer and assayed for viability using a Countess II automated cell counter (Thermo Fisher Scientific, Waltham, MA). From each sample a maximum volume containing 17,000 cells, having approximately >80% viable cells, were loaded onto the 10x Chromium (10x Genomics, Pleasanton, CA) controller for a targeted cell recovery of 10,000 cells (10x Genomics, Pleasanton, CA, Chromium Next GEM Single Cell 5’ v2 UserGuide RevD). cDNA amplification was carried out at 20 cycles of amplification for all samples and assayed for quality and concentration using a Bioanalyzer 2100 and High Sensitivity DNA Kit (Agilent, Santa Clara, CA).

### Ferret V(D)J enriched and 5’ gene expression library construction and sequencing

Construction of V(D)J libraries required the design and use of ferret-specific primers targeting the constant region of V(D)J transcripts for two subsequent enrichments as required by the 10x V(D)J workflow. An equimolar ratio of gene-specific primers was used to construct each pool, whereby Ig assays utilized a final concentration of 0.5 µM of each gene specific primer paired with 1µM 10x forward primer, and TCR assays utilized a final concentration of 1µM of each gene specific primer paired with 2µM of the 10x forward primer. Primer pools were constructed in volumes of 48uL to allow for direct substitution into the 10x workflow without further changes at this step. The number of amplification cycles was equalized between Ig and TCR enrichments such that each library was amplified for 12 cycles in the first enrichment and 10 cycles in the second enrichment. 5’GEX libraries underwent 16 cycles of final amplification. Otherwise, the protocol was followed according to the manufacturer’s specifications (10x Genomics, Pleasanton, CA, Chromium Next GEM Single Cell 5’ v2 UserGuide RevD).

Final V(D)J and 5’ GEX libraries were run on a Bioanalyzer 2100 with a High Sensitivity DNA kit (Agilent, Santa Clara, CA) to assess library size and concentration. Further analysis of library concentration was performed using a Qubit 3 fluorometer (Thermo Fisher Scientific, Waltham, MA) and combined with the sizing data from the Bioanalyzer to determine appropriate loading concentration for each library. Sequencing was performed using a NextSeq 500 sequencer (Illumina, San Diego, CA) utilizing 150 cycle kits in a paired end fashion. Runs were programmed to generate a 26bp read 1 sequence consisting of the 16bp 10x barcode and 10bp 10x UMI, two 10bp index reads, and a 122bp read 2 sequence of the cDNA insert. Sequencing depth was targeted at a minimum of 5,000 reads per cell for V(D)J libraries and 20,000 reads per cell for 5’ GEX libraries as recommended by the 10x workflow (10x Genomics, Pleasanton, CA, Chromium Next GEM Single Cell 5’ v2 UserGuide RevD).

### Single-cell immune repertoire sequencing analysis

Raw sequencing data were demultiplexed and converted to fastq files using the Cellranger (v7.0.0) command mkfastq. Next, raw sequencing reads were de novo assembled into contigs using the Cellranger vdj command. A de novo, rather than reference-based, contig assembly was performed due to the lack of a comprehensive collection of the ferret germline V(D)J sequences. To annotate the C- regions of the assembled V(D)J transcripts, we searched assembled contigs against inner-enrichment primers and the reference set of C-region sequences we compiled above, using the ublast (e value ≤ 1e−4) and usearch_global (sequence ID threshold = 0.9) commands, respectively, from the USEARCH (v10.0.240) ^46^ suite of sequence analysis tools. Subsequently, to annotate the V regions of the assembled V(D)J transcripts we again utilized the same custom IgBLAST to search all de novo assembled V(D)J contigs against the IMGT collection of V-region reference from ferret and related species, using the same protocol described for Iso-seq data analysis as described above. After V(D)J contig annotation, we systematically filtered all de novo assembled contigs as follows. First, we removed any contigs that represented misassembled chimeric transcripts (e.g., IGL primer alignment and IGHM C-region alignment) or off-target transcripts (e.g., primer match at the 3’ end but without a corresponding C-region alignment). Second, we removed contigs that did not meet the same filtering criteria we used in the annotation of Iso-seq reads to the IgBLAST database (variable gene hit with an e-value < -log_10_(50) and a joining gene hit < - log_10_(3)).

### Single-cell 5’ gene expression sequence analysis

Raw sequencing data were demultiplexed, aligned to the ferret reference assembly (GCA_011764305.2) with NCBI RefSeq Annotation release ID 102, and UMI-collapsed using cellranger (v7.0.0) commands mkfastq and count. We used the filtered feature-barcode matrices generated by cellranger, which contained only detected cell-associated barcodes. Cellranger used a cell-calling algorithm that was expected to identify populations of low RNA content cells and was based on the EmptyDrops method ^61^. UMI-count gene expression matrices were further processed using the protocol below using the Seurat package ^62^. Cells that appeared to be dead or dying were removed by filtering cell barcodes with greater than 5% of their UMIs mapping to any mitochondrial genes. Next, we removed technical doublets from our gene expression matrices using the doublet detection software Scrublet ^63^. We did not perform additional computational analysis to identify and correct for potential ambient RNA. Because the libraries for transcriptome and VDJ/VJ were prepared and sequenced separately, we reasoned that low quality cells by their transcriptome data might potentially contain valuable information about VDJ/VJ sequences and the diversity. For the PBMC sample, the minimum numbers of UMIs and genes of the remaining cell barcodes were 500 and 315. For the splenocyte sample, the minimum number of UMIs and genes of the remaining cell barcodes were 500 and 103.

We normalized the remaining gene expression data by log transforming the gene expression matrix using the “LogNormalize” method from Seurat. With this normalized matrix and canonical marker genes of cell cycle we used the CellCycleScoring function from Seurat to estimate cell cycle phase for each cell. The list of 97 human cell cycle gene symbols was from ^64^ and we used their ferret orthologs based on the ortholog gene groups publicly available from NCBI. Finally, we ran the normalization and scaling procedure SCTransform ^65^ with vars.to.regress equal to the S and G2M cell cycle scores assigned to each cell previously. Genes annotated with “C_region” and “V_segment” biotypes in the NCBI RefSeq Annotation were excluded from the list of 3,000 most variable genes in each dataset for the downstream analysis, to mitigate their potential interferences ^66^. The principal components analysis was performed using the normalized and scaled expression levels of the remaining most variable genes in each dataset. Based on Seurat’s recommendations, the first 30 principal components were used as input to uniform manifold approximation and projection (UMAP) and K-nearest neighbor cell clustering with a resolution parameter of 0.1 for both samples.

Cell type predictions of individual cells were performed using SingleR ^29^ using Seurat’s LogNormalized UMI counts and was independent of the cell clustering. We used the built-in reference Human Primary Cell Atlas ^67^ through the celldex package ^29^, and ‘pruned.labels’ for cell type prediction. Differential expression analysis of individual comparisons was performed using a Wilcoxon Rank Sum test within the FindMarkers function from Seurat with min.pct and logfc.threshold parameters, both set to the value of 0 to test for differential expression across all genes. Differentially expressed genes that had a p-value less than 0.05 were used for functional enrichment analysis using Ingenuity Pathway Analysis (Qiagen, CA). Separate tables of differentially expressed gene profiles of all cell clusters were generated using a Wilcoxon Rank Sum test within the FindAllMarkers function from Seurat with a min.pct parameter of 0.5 for all genes.

To further assess the B cell type predicted for cluster 4 in the splenocyte sample, from each cell we extracted RNA-seq reads aligned to the genomic regions annotated with ferret Ig variable genes. Because it is challenging to align individual RNA-seq reads to specific variable genes accurately, we chose to count the total number of unique reads aligned to each of the genomic regions (IGHV, IGKV, and IGLV) that we annotated Ig variable genes (regardless the specific locations that each read was aligned to), to approximate the detection of Ig transcript in each cell. To balance spurious alignments and potentially low efficiency of recovering VDJ transcripts directly from 5’ expression sequencing analysis, we filtered the obtained read counts using a minimum read count threshold. We then calculated the percentage of cells in each cell cluster with Ig heavy and light chain transcripts detected. The minimum read count thresholds were empirically determined by manually examining the expected differences in the obtained percentages between B and non-B cell clusters.

## Acknowledgements

Dr. Ed Breitschwerdt at North Carolina State University provided the ferret lymph node samples sequenced and analyzed in this study. Dr. Stacey Schultz-Cherry at St. Jude Children’s Research Hospital provided the ferret PBMCs used to validate the designed ferret-specific primers.

Thomas Rowe in Ted Ross’s lab at University of Georgia provided the ferret PBMCs and splenocytes used in the single cell sequencing analysis. Nedzad Music and Adrian Reber provided ferret splenocytes and helpful discussions. We thank Véronique Giudicelli for insightful scientific discussions and technical support concerning IMGT/HighV-QUEST analyses. We acknowledge interesting discussions with Corey Watson and the implication of Fratzeska Fragkiadi in TRA/TRD loci annotation.

This project has been funded in part with Federal funds from the National Institute of Allergy and Infectious Diseases, National Institutes of Health, Department of Health and Human Services, under Contract No. 75N93019C00052. We acknowledge the support of Immun4Cure IHU “Institute for innovative immunotherapies in autoimmune diseases” (France 2030/ANR-23-IHUA-0009) as well as the support of the Institut Universitaire de France. IMGT® is member of the French Infrastructure Institut Français de Bioinformatique (IFB) as well as member of BioCampus, MAbImprove and IBiSA.

## Author contributions

ESW – designed and performed experiments; analyzed data; wrote the manuscript

KY – designed and performed experiments; analyzed data; wrote the manuscript

TST – designed and performed experiments; analyzed data; wrote the manuscript

SS – designed and performed experiments; analyzed and curated data; wrote the manuscript

GZ – analyzed and curated data; wrote the manuscript

IS – analyzed and curated data; wrote the manuscript

GF – analyzed and curated data; wrote the manuscript

SK – analyzed and curated data; wrote the manuscript

NA – designed and performed experiments; analyzed and curated data; wrote the manuscript

HNB – designed and performed experiments; analyzed data; wrote the manuscript

HC – designed and performed experiments; analyzed data; wrote the manuscript

IAY – conceptualized and supervised the project; designed experiments; analyzed data; wrote the manuscript

XP – conceptualized and supervised the project; designed experiments; analyzed data; wrote the manuscript

## Declaration of interests

X.P. is the Founder and CEO and has an equity interest in Depict Bio, LLC. The terms of this arrangement have been reviewed and approved by NC State University in accordance with its policy on objectivity in research.

The findings and conclusions in this report are those of the authors and do not necessarily represent the official position of the Centers for Disease Control and Prevention.

**Inclusion and diversity statement (optional)**

## Additional supplemental tables

Table S1: Summary of the ferret Iso-Seq data and the abundance of ferret C-region isoforms (excel file).

Table S2: Ferret Ig and TCR C-region cDNA consensus sequences, annotations and description of allelic sites, and V, D, J gene annotations on the ferret reference genome assembly (excel file). The “IGHV Provisional” tab provides provisional names for IG V genes found on genomic contigs not linked with constant regions. The “Allelic Variants” tab describes allelic variants of IG and TR constant regions identified in transcriptomes and genomic sequences, including CCS counts for each allele. Transcriptomes from splenocytes from two (SRR29376976), four (SRR29376975), three (SRR29376974), or two (SRR26825671 and SRR26825672) were screened for the number of exact matches to the sequences as shown. Only those genes for which allelic variants were identified are shown here. Transcriptome datasets are available from BioSamples SAMN41805205, SAMN41805206, SAMN41805207, SAMN38182219 and SAMN38182220. Ferret Ig and TCR C-region annotations on the 10 additional ferret genome assemblies for the *in silico* PCR analysis of ferret-specific Ig and TCR primers are available on GitHub.

Table S3: Ferret Ig and TCR C-region specific primer sequences and the comparison of V(D)J contigs with primer hit and with vs. without the corresponding C-region match (excel file).

Table S4. Differentially expressed genes for each cell cluster of the ferret PBMC and splenocyte samples (excel file).

Table S5: Comparison of ferret cell type predictions using human references (in the same Supplementary Data file with Supplementary Figures).

Table S6: Functional enrichment analysis of differentially expressed genes between selected cell clusters identified in the ferret PBMC and Splenocyte samples (excel file).

